# CRISPR-based knockin mutagenesis of the pioneer transcription factor FOXA1; optimization of strategies for multi-allelic proteins in cancer cells

**DOI:** 10.1101/2020.10.27.356824

**Authors:** Shen Li, Joseph P. Garay, Colby A. Tubbs, Hector L. Franco

## Abstract

Precise genome engineering of living cells has been revolutionized by the introduction of the highly specific and easily programmable properties of CRISPR-Cas9 technology. This has greatly accelerated research into human health and has facilitated the discovery of novel therapeutics. CRISPR-Cas9 is most widely employed for its ability to inactivate, or knockout, specific genes, but can be also used to introduce subtle site-specific substitutions of DNA sequences that can lead to changes in the amino acid composition of proteins. Despite the proven success of CRISPR-based knockin strategies of genes in typical diploid cells (i.e. cells containing two sets of chromosomes), precise editing of cancer cells, that typically have unstable genomes and multiple copies of chromosomes, is more challenging and not adequately addressed in the literature. Herein we detail our methodology for replacing endogenous proteins with intended knockin mutants in polyploid cancer cells and discuss our experimental design, screening strategy, and facile allele-frequency estimation methodology. As proof of principle, we performed genome editing of specific amino acids within the pioneer transcription factor FOXA1, a critical component of estrogen and androgen receptor signaling, in MCF-7 breast cancer cells. We confirm proper levels of mutant FOXA1 protein expression and intended amino acids substitutions via western blotting and mass spectrometry. In addition, we show that mutant allele-frequency estimation is easily achieved by TOPO cloning combined with allele-specific PCR, which we later confirmed by next-generation RNA-sequencing. Typically, there are 4 - 5 copies (alleles) of FOXA1 in breast cancer cells making the editing of this protein inherently challenging. As a result, most studies that focus on FOXA1 mutants rely on ectopic overexpression of FOXA1 from a plasmid. Therefore, we provide an optimized methodology for replacing endogenous wildtype FOXA1 with precise knockin mutants to enable the systematic analysis of its molecular mechanisms within the appropriate physiological context.

## Introduction

CRISPR (clustered regularly interspaced short palindromic repeats) and its associated components was originally discovered as an RNA-guided adaptive immune system in bacteria and archaea (1-3). This defense mechanism allows microorganisms to integrate short fragments of exogenous, pathogenic DNA sequences into their own genome to generate a library of CRISPR RNAs (crRNAs) that serve as a memory of past genetic aggressions. Once transcribed, these crRNAs form a complex with endogenous trans-activating crRNAs (tracrRNA) and nucleases called Cas proteins to form active ribonucleoprotein complexes that can search and destroy foreign invading DNA sequences (3-8). The programmable properties of CRISPR that originated as a natural defense mechanism in bacteria has now been repurposed for RNA-guided cleavage of any DNA sequence in mammalian cells (3, 9).

The Type II CRISPR-Cas9 system has been engineered for optimal efficiency in human cells. This technology combines a crRNA and tracrRNA into a single-guide RNA (sgRNA) that can be programmed to deliver the Cas9 nuclease to any specific DNA sequence (9, 10). Once Cas9 is bound to a target sequence, it creates a double stranded break (DSB) inducing cellular pathways needed for DSB repair. It is at this point that researchers can edit the sequences in the vicinity of the DSB. There are two main DNA repair pathways in mammalian cells, Non-Homologous End Joining (NHEJ) repair pathway and the Homology Directed Repair (HDR) pathway (11-13). NHEJ is typically the choice for repair because it is less energetically demanding than HDR and it does not require a repair template. However, NHEJ is more error-prone and can produce frameshift mutations that abrogate gene function. This approach allows researchers to study the consequences of the ablation of a specific DNA sequence or gene and, in most cases, creating knockout cell lines is relatively straightforward (9). Alternatively, the DSB may be repaired by the higher fidelity Homology Directed Repair (HDR) pathway with the aid of a repair template. This repair template can be delivered exogenously and its sequence can be designed to contain specific nucleotide substitutions that can be incorporated into the endogenous DNA target sequence. If the desired nucleotide substitutions are made within the coding sequences of proteins, researchers have the ability to change the resulting amino acid composition of the target protein (Fig. 1). These site-specific amino acid substitutions are referred to as Knockin (KI) mutations (9, 14-16). As compared to typical Knockout experiments, the success of CRISPR mediated Knockin experiments depend on both the efficiency of DSB caused by the CRISPR-Cas9 and the effective delivery of the repair template. Of note, some cancer cell lines have inactivated HDR pathways and can only repair double stranded breaks using the error prone NHEJ pathway effectively nullifying the ability to create knockins in that cell line. These considerations make knockins considerably more challenging than knockouts. In most cases, especially in polyploid cells, this system is not entirely efficient and successfully edited KI alleles typically coexist with either wild type or frameshift alleles. This illustrates an inherent challenge in applying CRISPR directed site-specific mutagenesis to multi-allelic genes in polyploid cancer cell lines; a scenario that is likely common and of importance to the scientific community.

**Figure 1.**
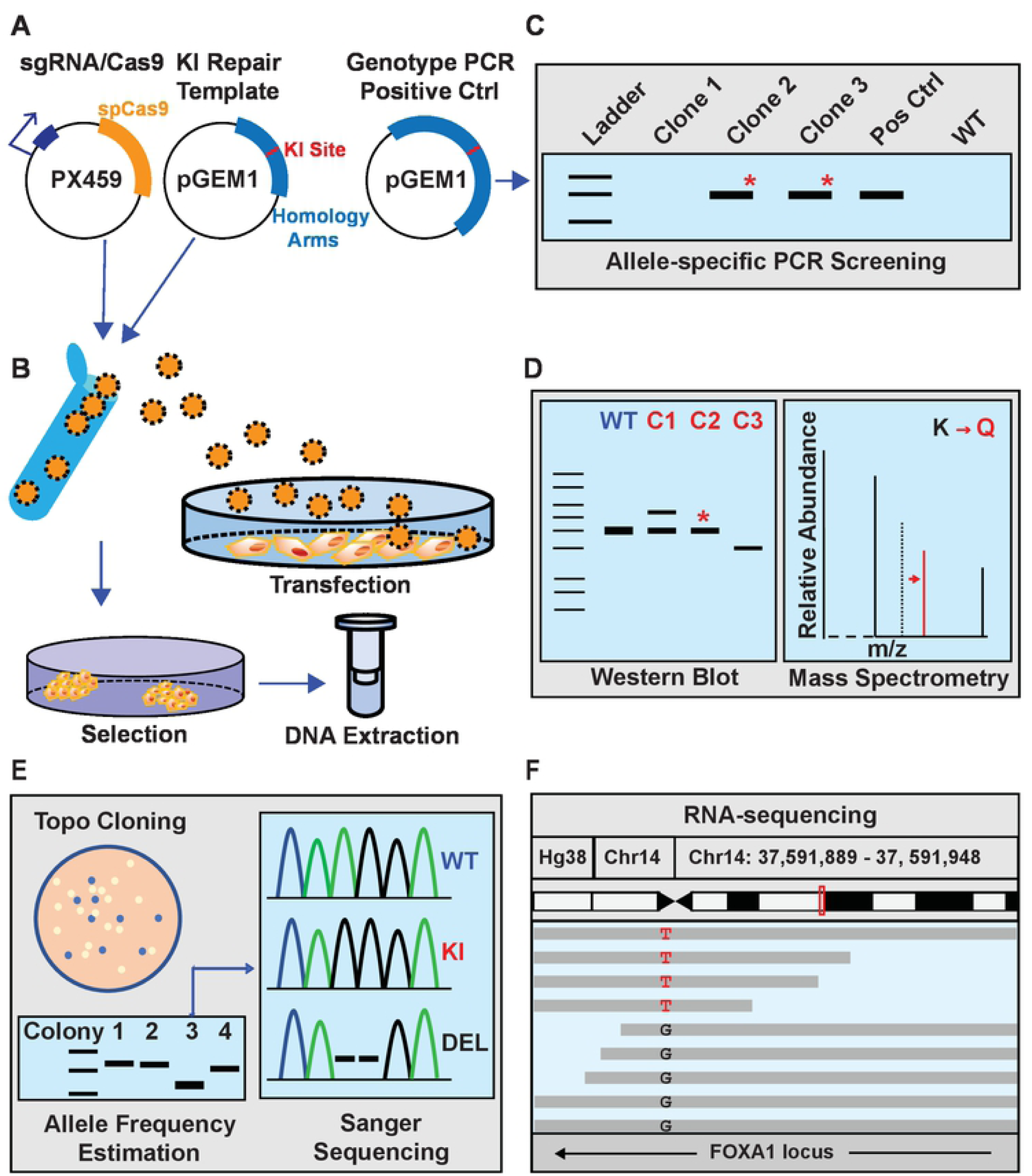
General workflow for CRISPR-based site-specific mutagenesis of multi-allelic genes in cancer cell lines. **A**. Three plasmids used for mutagenesis. The plasmid expressing spCas9 and the sgRNA (pX459) and the repair template plasmid containing the homology arms to the intended target site as well as the desired nucleotide substitutions for mutagenesis (pGEM1) are transfected into the cells. A third plasmid, containing a larger fragment of the endogenous locus sequence plus the desired nucleotide substitutions for mutagenesis is used as a positive control for the allele-specific PCR genotyping strategy. **B**. Transfections, antibiotic selection, colony picking, and genomic DNA extraction for genotyping of potential knockin clones is discussed in the text. **C**. Allele-specific PCR is used as a screening strategy to identify clones that have successfully integrated the repair template sequence with desired nucleotide substitutions (knockin). **D**. Western blotting and mass spectrometry (LC-MS/MS) is used to confirm that knockin clones successfully produce full length knockin proteins. **E and F**. Allele frequency estimation is performed using TOPO cloning-based screening of genomic DNA extracted from individual knockin clones and/or the use of next-generation sequencing approaches such as RNA-sequencing to measure the abundance of RNA transcripts that contain the desired knockin mutations. The details of each step are discussed throughout the text.

For our studies, we sought to specifically modify the pioneer transcription factor FOXA1 in the Luminal A breast cancer cell line MCF-7. Pioneer factors are characterized for their ability to associate with condensed chromatin independently of other factors to directly modulate chromatin accessibility. FOXA1 is known as a key pioneer transcription factor for nuclear receptors such as estrogen receptor (ER) and androgen receptor (AR) in breast and prostate cancers respectively (17-19). FOXA1’s pioneering functions stem from its protein structure that possess features of both linker histones and conventional transcription factors allowing it to displace linker histones in compacted nucleosomes and allow access to other transcription factors such as ER and AR (17-19). Thus, altering the amino acid composition of FOXA1 will alter its pioneering function across the genome and consequently alter ER and AR signaling pathways. Of note, there are multiple copies of FOXA1 in breast cancer cells (typically 4 −5 alleles) making the editing of this protein inherently challenging. As a result, virtually all publications that focus on FOXA1 mutants rely on ectopic overexpression of FOXA1 from a plasmid. CRISPR-based mutagenesis of FOXA1 at the endogenous locus offers several advantages over the more traditional approach of ectopic overexpression from a plasmid. First, it is possible to eliminate the wild-type version of the protein by replacing all of the copies with the intended knockin variants. This provides a homogenous model system where the phenotypic outcomes are solely due to the knockin variants and not due to a combination of knockin and wild-type proteins typically seen with ectopic overexpression experiments. Second, editing directly within the endogenous loci will maintain the local chromatin and regulatory mechanisms intact, thus producing physiologically relevant levels of protein expression. Finally, the CRISPR-Cas9 system is more easily implemented while introducing similar levels or sometimes less off-target effects as compared to other genomic engineering techniques such as Zing-finger nucleases (20, 21) and TALENs (22-24).

Herein, we have detailed our strategy for editing FOXA1 in the polyploid cancer cell line MCF-7. We provide a reproducible protocol, screening methodology, and allele-estimation strategy, as well as verification steps and important considerations for characterizing KI cell lines. Figure 1 illustrates the general workflow and we have outlined our procedure in Table 1.

**Table 1:** Tasks and timeline for CRISPR-Cas9 based site specific mutagenesis in polyploid cancer cells.

| Task |  | Duration |
| --- | --- | --- |
| 1 | Guide sgRNA design and Cas9/sgRNA plasmid construction | 1 week |
| 2 | Design of the HDR template and KI screening control plasmid | 2 weeks |
| 3 | Cell transfection, selection, collection, and DNA purification | 8 weeks |
| 4 | Genotyping through allele-specific PCR | 2 weeks |
| 5 | Protein expression and mass spectrometry of Knockin clones | 2 weeks |
| 6 | Allele frequency estimation in knockin clones | 2-4 weeks |

## Materials and Methods

### sgRNA design and Cas9/sgRNA plasmid construction

One of the most critical steps in proper editing of mammalian cancer cell lines is the design of the guide RNA (henceforth referred to as *sgRNA*) needed to deliver Cas9 to your genomic region of interest. For this application we used the mammalian expression vector pSpCas9(BB)-2A-Puro (pX459) V2.0 that transcribes the sgRNA from a U6 promoter, produces Cas9 derived from *S. pyogenes* (spCas9), and contains a puromycin resistance gene needed for clonal selection (Fig.1). This plasmid was generated by Feng Zhang’s lab and is available from the plasmid repository Addgene (plasmid ID 62988) (9).

For our proof of principal example, we focused on site-specific amino acid substitutions of the pioneer transcription factor FOXA1 in the breast cancer cell line MCF-7. There are 4-5 copies of FOXA1 in MCF-7 cells depending on the strain and source of MCF-7 cells used. We specifically targeted lysine 295 (K295) and detail the steps to mutate this lysine residue to glutamine (K295Q) (Fig.1).

There are several online resources available for sgRNA design. CRISPOR (http://crispor.org/) is a easy to use, freely available, and a well-vetted design tool that generates several putative guide RNA target sequences along with a series of *specificity, efficiency*, and *off-target* scores that can be used to select the best sgRNA (25). The human genomic sequence of FOXA1 was obtained from the National Center for Biotechnology Information database (NCBI, FOXA1: NG_033028.1) and 300bp of genomic sequence, centered on the knockin (KI) site of interest (Lysine K295), was submitted to the CRISPOR website. sgRNA sequences with a value of 80 or higher for the *MIT* score and 50 for the *Doench* score were considered good candidates. To minimize off-target effects, sgRNAs that were predicted to have few or no off-target sites were selected. Of note, sgRNA sequences depend on the nature of the genomic sequence at the target loci, and in some cases high specificity scores and efficiency scores are simply not obtainable. If all sgRNAs do indeed show potential off-target sites, then preference is given to sgRNAs whose off-target sites that are located in non-coding regions versus coding regions of the genome. The targeting sequence for K295 of FOXA1 is shown below. This includes the 20bp targeting guide sequence (blue font) plus 3bp for the PAM sequence (orange font). The PAM (protospacer adjacent motif) is a short sequence that must immediately follow on the 3’ end of the targeting guide sequence. These 3bp are required for proper cleavage by the Cas9 nuclease. The PAM requirement for *S. pyogenes* Cas9 nuclease is any nucleotide followed by two guanines (5’-NGG)(9).

> FOXA1 K295 targeting sequence: 5’-AGCGGGGGCAGCGGCGCCAAGGG-3’

To prepare oligonucleotide sequences for cloning into the pX459 plasmid, the 3’ NGG PAM sequence was deleted from the CRISPOR output sequence, and the BbsI restriction enzyme motif sequence 5’-*CACC-3’* was added to the 5’ end (green font) of the remaining 20 bp targeting guide sequence (blue font) for cloning into the pX459 plasmid. Due to the nature of the U6 promoter from which the sgRNA is transcribed, a guanine (G) is added immediately 5’ of the oligonucleotide in order to allow for efficient transcription (black font) (9). This generates the forward sgRNA oligonucleotide:

> FOXA1 K295 Forward sgRNA Oligo: 5’-CACCGAGCGGGGGCAGCGGCGCCAA-3’

To generate the reverse strand oligo, the reverse complement was obtained from the 20 bp targeting guide sequence (blue font) and the BbsI restriction enzyme motif sequence 5’-*AAAC-3’* was added to the 5’ end (green font). This completes the reverse strand sgRNA oligonucleotide:

> FOXA1 K295 Reverse sgRNA Oligo: 5’-AAACTTGGCGCCGCTGCCCCCGCTC-3’

As with most CRISPR-based experiments, it is always advisable to design more than one sgRNA. Depending on the cell context, some sgRNAs will work better than others. Specifically, when dealing with gene editing of multi allelic loci, more than one sgRNA is needed in the case where some, but not all, of the alleles were successfully edited. In most cases, the unedited loci will contain small indels that will ablate the binding site or efficacy of the sgRNA. Therefore, a second sgRNA targeting a different location (slightly upstream or downstream) of the first sgRNA will allow for a second round of transfection to target the un-edited alleles and create homogenous clones. For FOXA1, we designed three different sgRNAs, all centered at the target amino acid (K295) (Fig. 2). For efficient editing it is recommended to design the sgRNAs at or immediately adjacent to the desired target amino acid.

> FOXA1 K295 sgRNA #1 5’-TGGGGTTAGAGGCGCCAGAG-’3
>
> FOXA1 K295 sgRNA #2 5’-AGCGGGGGCAGCGGCGCCAA-3’
>
> FOXA1 K295 sgRNA #3 5’-AGGGGTCCTTGCGGCTCTCA-3’

**Figure 2.**
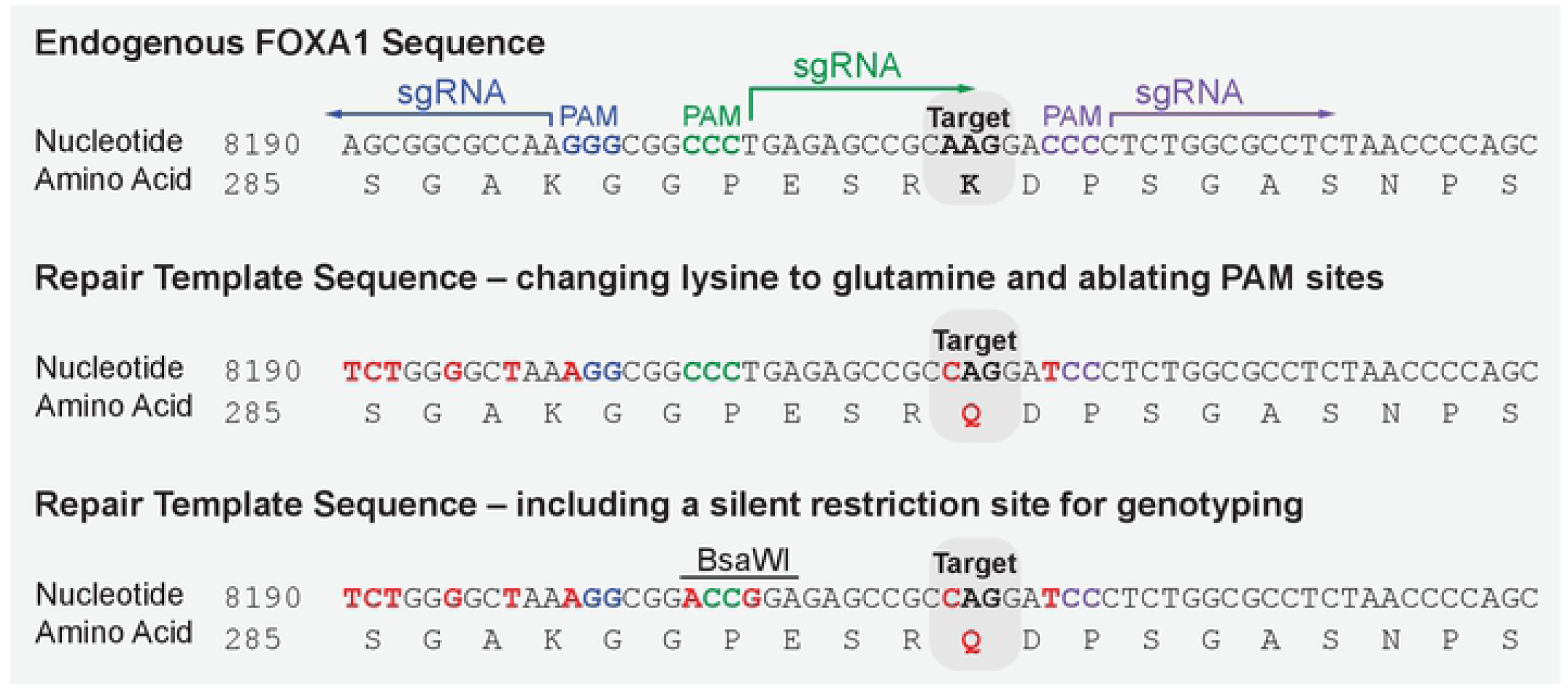
Preparation of the repair template sequence for precise genome editing. The human FOXA1 reference sequence is shown at the top. The target site lysine 295 (K295) is highlighted and the three sgRNAs are shown along with their associated PAM sites. To design the repair template, the desired nucleotide substitutions are made to specifically change the amino acid sequence of the target protein. In addition, several silent mutations are introduced to alter the endogenous PAM sites to avoid repeated cleavage by Cas9 (middle sequence). Silent mutations that created a restriction enzyme recognition site were also designed into the repair template to provide an alternative genotyping strategy (bottom sequence).

For each of the three guide sequences, the BbsI restriction enzyme motifs were added to the 5’ and 3’ ends (as shown above) and the resulting oligonucleotides were ordered from Integrated DNA technologies (IDT). The sgRNA oligonucleotides were resuspended in water to a final concentration of 100 uM and the corresponding forward and reverse oligos were annealed and cloned into the pX459 plasmid using the protocol described by Ran F.A. *et. al*. (2013) *Nat. Prot* (9). Briefly, 1 ul of the forward sgRNA oligo (100 uM) was annealed with 1ul of the reverse sgRNA (100 uM) using 1 ul 10X T4 ligation buffer (New England Biolabs, NEB), 1 ul T4 PNK (NEB), and 6 ul of water. The annealing reaction was performed in a thermocycler using the following parameters: 37°C for 30 minutes, 95°C for 5 minutes, and final ramp down to 25°C at 5°C per minute (about 15-minute ramp down).

The annealed sgRNA oligos were diluted 1:200 in water and ligated into the pX459 plasmid that has been linearized with restriction enzyme BbsI (New England Biolabs). The ligation mixture contained 2 ul of diluted sgRNA oligos, 100 ng of linearized PX459 plasmid, 2 ul of 10X T4 ligation buffer, and 1 ul T4 DNA ligase (NEB). The ligation reaction was performed in a thermocycler at 37°C for 5 min followed by 21°C for 5 min.

The ligated plasmid was transformed into DH5α competent E. coli cells (Invitrogen, 18265017) and spread onto LB agar plates containing 100 ug/ml ampicillin. Plasmids were prepared using silica-membrane based kit (Plasmid Miniprep Kit, Qiagen) following the manufacturer’s instructions and quantified using Nanodrop Spectrophotometer. Successful integration of the sgRNA was confirmed by Sanger Sequencing using the common “U6” sequencing primer, 5’-GGCCTATTTCCCATGATTCC-3’.

### Preparation of the repair template sequence for precise genome editing

There are several key features of the repair template that need to be considered when designing the sequences for site-specific mutagenesis. The repair template should include the desired nucleotide substitutions intended to specifically change the amino acid sequence of the target protein. In addition, the repair template should include silent mutations (i.e. nucleotide substitutions that do not change the amino acid sequence) designed to alter the endogenous PAM site or the endogenous 20bp gRNA targeting sequence in order to avoid repeated cleavage by Cas9 once the cell has successfully repaired the DSB (Fig 2). Silent mutations that create a restriction enzyme recognition site may also be designed into the repair template in order to provide an alternative genotyping strategy if desired. This must be designed in a way that does not alter the amino acid sequence of your protein or the open reading frame.

To design the repair template for modification of K295 on FOXA1, the entire genomic sequence (including the 5’ UTR, introns, and 3’UTR) was obtained using the NCBI genomic database (FOXA1: NG_033028.1). The genomic sequence was annotated to identify the start codon, open reading frame, and corresponding amino acids. The PAM recognition sites for each of the three sgRNAs are also annotated in the sequence (Fig 2).

To generate the intended mutation, the repair template sequence was designed to contain a single nucleotide substitution needed to convert lysine at position 295 into a glutamine (AAG to CAG). In addition, silent mutations were introduced into the repair template sequence in order to ablate the PAM recognition sequence (or other significant nucleotides within the sgRNA binding site) (26). This prevents additional rounds of cutting by Cas9 once the DSB has been successfully repaired using the provided repair template (Fig. 2). As a final step, we introduced silent mutations that created a restriction enzyme site within the target locus using the Watcut online tool (http://watcut.uwaterloo.ca/). This provides an alternative genotyping strategy (genomic PCR followed by restriction digest) to identify clones that have successfully repaired the endogenous locus using the repair template. For FOXA1, the restriction enzyme BsaW1 was used because there are no naturally occurring restriction sites within 300bp upstream or downstream of the edited region (Fig. 2). For genotyping, primers were designed to amplify 300bp upstream and 300bp downstream of the target amino acid and thus limited our restriction enzyme motif search to this stretch of DNA. Genotyping strategies are discussed in more detail below.

### Preparation of the repair template plasmid and the positive control plasmid needed for allele specific PCR screening

Once the sequence for the repair template has been designed, the repair plasmid may be built using simple cloning plasmid backbones (smaller plasmid backbones are preferred). The entire plasmid, including the repair sequence can be custom ordered from most nucleic acid companies or built using traditional plasmid editing techniques such as site-directed-mutagenesis (SDM). For FOXA1, we generated two different plasmids, the repair template plasmid and a screening-PCR positive control plasmid to be used to validate our allele-specific PCR (Fig. 3).

**Figure 3.**
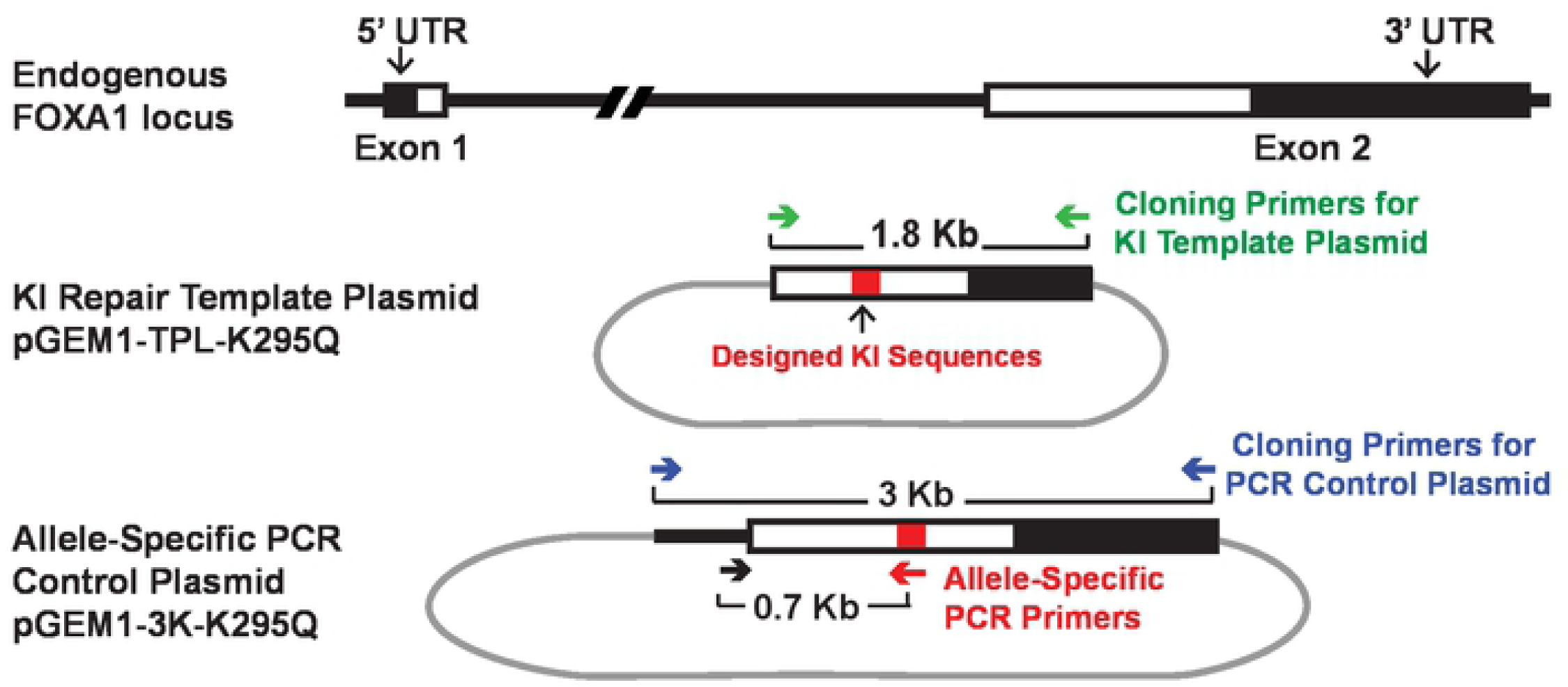
Repair template plasmid and PCR screening positive control plasmid preparation. The human FOXA1 endogenous locus is shown at the top. Two different plasmids were generated, the repair template plasmid and an allele-specific PCR positive control plasmid to be used during the screening of potential KI clones. The repair template plasmid was built on the pGEM1 backbone and contains FOXA1 repair template sequence (shown in Figure 2) along with FOXA1 homology arms of 600bp upstream and 1200bp downstream of K295. The allele-specific PCR positive control plasmid was intentionally designed to contain more of the endogenous sequence of FOXA1 as well as the repair template sequence to be able to design our allele-specific PCR. The forward primer only hybridizes to this plasmid and not the repair template plasmid thus allowing us to detect successful KI editing within the endogenous locus of FOXA1 and not random integration of the repair plasmid.

To create the repair template plasmid, we used the following primers *FOXA1KITPL-F:* 5’-CAAGCTTGCCATGAACAGCATGACTG-3’ and *FOXA1KITPL-R:* 5-CGGATCCCTGAGAAGCAAATGGCTCTG-3’ to amplify a 1.8kb region surrounding our FOXA1 K295 site from MCF-7 genomic DNA (Fig. 3). The Phusion High Fidelity DNA Polymerase (NEB Cat # M0530L) was used per manufacturer’s instructions in order to ensure proper amplification of the large 1.8 kb fragment. The primers were designed with restriction enzyme site overhangs (BamHI and HindIII) to facilitate cloning into the pGEM-1 plasmid (Promega) (Fig. 3). Of note, the KI site (K295) divides the 1.8 kb genomic fragment into 600 bp and 1200 bp homology arms, and the design for this particular locus ensures both targeting and screening efficiency (27, 28). Homology arm length can vary depending on the target locus from less than 50bp on either end to over 1000bp (29, 30). Once cloned into the pGEM1 plasmid, we used the Q5 Site-Directed mutagenesis Kit (NEB Cat# E0554S) per manufacturer’s instructions to introduce the specific nucleic acid substitutions needed to create the repair template sequence shown in Figure 2. The primers used for site-directed mutagenesis were *SDM K295Q F:* 5’-CCGGAGAGCCGCCAGGATCCCTCTGGCGCCTCTAACCC-3’ and *SDM K295RQ R:* 5’-TCCGCCTTTAGCCCCAGAGCCCCCGCTTCCGCTCCC-3’. Once completed, the repair template plasmid was prepped using silica-membrane based kit (Plasmid Miniprep Kit, Qiagen) and sent off for Sanger Sequencing to confirm proper assembly of the plasmid.

To create the plasmid that will be used as a positive control for the PCR reaction used for genotyping, the following primers *FOXA1E2TPL-F:* 5’-CAAGCTTTTGACAAACTGTGTCACC-3’ and *FOXA1E2TPL-R:* 5’-CGGATCCACCCGTCTGGCTATACTAAC-3’ were used to amplify 3kb region surrounding the FOXA1 K295 site from MCF-7 genomic DNA. As done with the repair template primers, these primers were also designed with restriction enzyme site overhangs (BamHI and HindIII) to facilitate cloning into the pGEM-1 plasmid (Promega) (Fig. 3). The final step was to introduce the repair template nucleotide substitution sequences into this plasmid. Again, the Q5 Site-Directed mutagenesis Kit was used along with the same SDM primers. This plasmid was intentionally designed to contain more of the endogenous sequence of FOXA1 as well as the repair template sequence to be able to design our allele-specific PCR (Fig. 3). This is discussed further below. After plasmid prep and Sanger sequencing, we now have a completed allele-specific screening-PCR positive control plasmid.

### Transfection and selection of knockin cell lines

There are many methods for delivering the sgRNA/Cas9 and the repair template plasmid (i.e. electroporation, viral delivery, or lipid-based transfections). Each strategy has its own advantages and disadvantages and have been well discussed in these reviews (31, 32). It is important to choose a transfection method that will be suitable with the intended cell line. We recommend optimizing this procedure with plasmids expressing fluorescent reporters to visualize transfection efficiency and monitor cell toxicity. For MCF-7 breast cancer cells, we used the lipid-based Lipofectamine 3000 (Thermo Fisher) transfection method due to its low cost and feasibility. If a viral delivery method is preferred, these plasmids would need to be reconstructed accordingly. In addition, the repair template can be delivered into the cell in a variety of forms. For example, the repair template can be provided to the cell as a circular plasmid, a linearized plasmid, or an isolated template sequence lacking a plasmid backbone; all have been shown to successfully generate KI cell lines. After testing these options, we found the circular plasmid to be the most efficient and reliable method to deliver a repair template sequence into MCF-7 cells.

For transfection, MCF-7 cell lines are cultured in DMEM containing 10% FBS and 1% Pen-Strep and kept in an incubator at 37°C with 5% CO2. 24 hours prior to transfection, 80,000 cells were seeded in each well of a 12-well cell culture plate. Optimal seeding density for each cell line should be empirically determined to avoid premature confluency after 4 days of cell culture. The media was replaced with 1 mL of antibiotic-free culture medium 1 hour prior to transfection. Lipofectamine 3000 was used to transfect the cells following the manufactures instructions. Briefly, the transfection complexes contained 1.5 ul of Lipofectamine, 500 ng of pX459 plasmid (sgRNA/Cas9), 500 ng of the repair template plasmid, and 2 ul of P3000 in a total volume of 100 ul of Opti-Mem reduced serum media (Thermo Fisher). After an 8-hour incubation, the cell media/transfection complexes were replaced with fresh antibiotic-free media to avoid cell toxicity. The puromycin antibiotic was used for selection of cells that had been successfully transfected. Cells were grown in complete media containing 1 ug/mL puromycin for 5-7 days, replacing the culture medium daily. Halfway through selection, the cells were trypsinized and transferred to 60 mm dishes to spread the remaining cells out and allow for individual clones to grow without colliding into each other. A batch of un-transfected cells were used to determine when selection with puromycin was complete (i.e. when complete cell death was observed in the control dish).

There are many strategies to pick clones after puromycin selection (serial dilutions, cell lifting, FACS sorting, etc) (9). Once visible colonies appeared on the plate, we used glass cloning rings (fixed to the plate with sterile Vaseline) to isolate a particular clone inside the glass ring. Then, trypsin was added to the glass ring to detach the cells and transfer the isolated clone to a separate well within a 24-well cell culture plate. After expansion of the clone to confluency, the cells were trypsinized and split evenly into two different plates: one plate for genomic DNA extraction and genotyping, and the other plate for maintenance of the clone for continuous culture.

## Results

### Genotyping clonal cell populations using allele-specific PCR

We sought to develop a robust genotyping strategy to screen our transfected cell lines for our desired knock-in allele. To do this, we used allele-specific PCR. This PCR technique relies on the presence of the nucleotide substitutions specified by the repair template within the endogenous FOXA1 locus. We designed an allele-specific PCR primer that only anneals to the repair template sequence (Fig. 3). This primer sequence needs to be sufficiently different than the wild type DNA sequence (include 3 to 4 mismatches relative to wild type sequence). This will help avoid false positives during PCR screening. This allele-specific primer was paired with a second primer that is sufficiently upstream of the knockin site that it cannot anneal anywhere within the 5’ homology arm sequence included in the repair template. This generates a specific DNA product that is only possible if there is a successful knockin at the endogenous locus (Fig. 4). Importantly, the primer that pairs with the allele-specific primer must be within the genomic region that is included in the screening-PCR positive control plasmid but not the repair template plasmid (i.e. the forward primer should not be able to anneal to the repair template plasmid) (Fig. 3). This ensures that the allele-specific PCR screen detects only true knockin events and not random integration of the repair template sequence in the host genome. The ASP primers used for FOXA1 K295Q were *K295Q ASP FW:* 5’ ACATGTCCTATGCCAACCCG 3’ and *K295Q ASP RV:* 5’ GCGCCAGAGGGATCCTG 3’.

**Figure 4.**
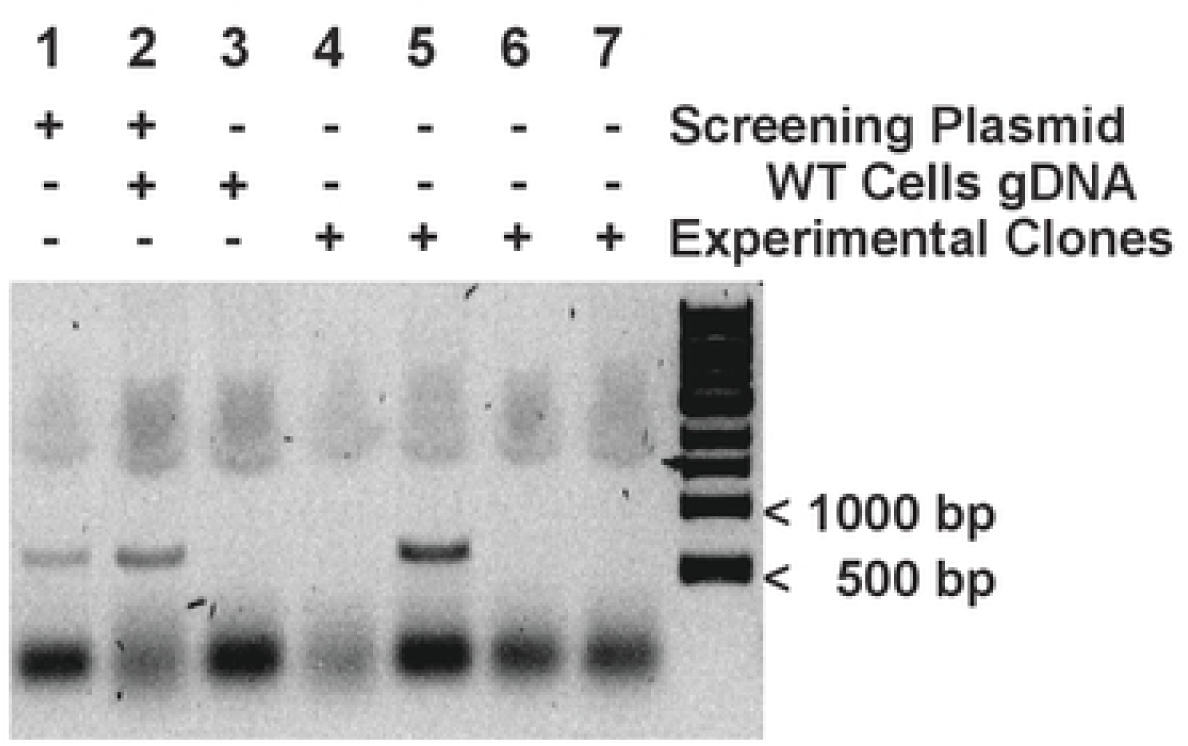
Allele-specific PCR reveals successful FOXA1 knockin clones. Agarose gel electrophoresis of allele-specific PCR results from puromycin resistant MCF-7 clones showing successful editing of FOXA1 at K295 (lane 5). Lanes 1 and 2 are positive controls for the PCR using the allele-specific PCR positive control plasmid shown in Figure 3. Genomic DNA from wild-type MCF-7 cells was used as a negative control.

For genotyping of our clonal cell lines for successful knockins, we used 200ng of purified genomic DNA for template in the PCR reaction. Genomic DNA was extracted from the individual clones using the Genomic DNA Extraction Kit (Qiagen). As a positive control for the PCR reaction, we mixed 0.4 pg of the screening-PCR positive control plasmid with 200ng of genomic DNA to mimic physiological concentrations of this sequence. Based on the plasmid size and polyploid nature of the MCF7 cell line, we determined that 0.4 pg of this plasmid has the same amount of FOXA1 alleles (3-5) as that in 200 ng of genomic DNA (33). As a negative control, 200 ng of wildtype genomic DNA was used. Thus, successful allele-specific PCR would generate a 650bp PCR product in clones that had successful knockin events but not in wildtype/unedited genomic DNA alone (Fig. 4).

For the PCR reaction the DNA Taq Polymerase with ThermoPol Buffer from New England Biolabs (NEB, M0267L) was used per manufacturer’s instructions. The thermocycling conditions were; initial denaturation at 95oC for 1 minute followed by 24 cycles of 95oC for 15 seconds followed by annealing step at 68oC for 15 seconds and extension at 68oC for 42 seconds. After every 3 cycles, the annealing temperature is decreased by 2 degrees until reaching a temperature of 54oC. At 54oC, five final cycles are completed (for a total of 29 cycles). The final extension is at 68oC for 5 minutes and the sample is held indefinitely at 4oC. PCR products were then electrophoresed at 120 volts for 45 minutes on a 0.8% agarose gel stained with SYBR Safe nucleic acid stain (Thermo Fisher) (Fig. 4).

### Knockin allele frequency estimation using TOPO cloning, allele-specific colony PCR, and RNA-sequencing

For all CRISPR-based genome engineering experiments, it is important to estimate the allele frequency of successful knockin events, especially when working with polypoid cell lines. We used Topoisomerase I based cloning (TOPO TA Subcloning kit, Thermo Fisher) followed by colony PCR screening to examine the allele composition and knockin frequency of FOXA1 in our edited MCF-7 cells. We reasoned that by amplifying the genomic DNA surrounding the KI site we would capture all of the different editing events that occurred across the multiple alleles within an individual clonal cell line. By designing primers that are well outside the KI site, the resulting PCR product would contain properly edited alleles together with any remaining indel or wildtype alleles. Subsequent cloning of the entire heterogenous PCR product into a TOPO vector (which allows for simple cloning of PCR products without using restriction enzymes), transformation into bacteria, and Sanger sequencing of the resulting bacterial colonies would allow for the estimation of KI alleles in a given cell line based on the ratio of KI alleles to wild type/indel alleles found (Fig. 5).

**Figure 5.**
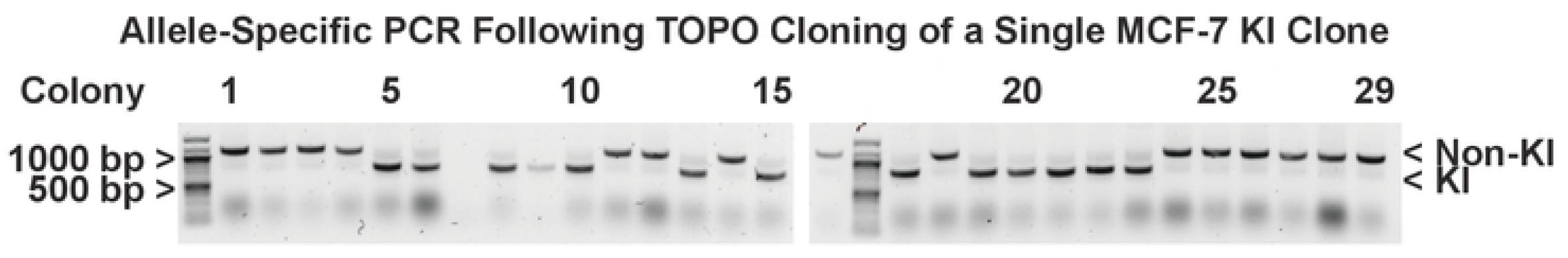
Knockin allele frequency estimation using TOPO cloning coupled to allele-specific colony PCR. Allele-specific colony PCR results from a single KI clonal cell line where 30 bacterial colonies resulting from TOPO cloning of the endogenous FOXA1 target sequence (K295) were screened. Successful knockin alleles give a 700bp product and wildtype alleles give a 1000 bp product. This experiment shows that of the 30 bacterial colonies screened for this one cell line, 13 colonies contained the KI allele, suggesting that about one third of the FOXA1 alleles were successfully edited.

To estimate the KI frequency of FOXA1, genomic DNA was purified from each of clonal cell lines that showed a positive KI result from genotyping. A 1 kb region surrounding the FOXA1 KI site was amplified using the following primers *K295Q ASP FW:* 5’-ACATGTCCTATGCCAACCCG-3’ and *F1K SEQ R:* 5’-GTGCAGCTGGGACTCGTGGG-3’. The PCR reaction was performed with Taq DNA Polymerase (NEB Cat # M0273S) and, after agarose gel electrophoresis and gel purification, the 1 kb PCR product was ligated into the TOPO 2.1 vector. A total of 5 ul of the ligation reaction was transformed into DH5a competent cells and 100 ul of the reaction was spread on LB-ampicillin plates. There are between 3-5 alleles of FOXA1 in MCF-7 cells, therefore we reasoned that screening about 30 bacterial colonies resulting from the TOPO cloning reaction would allow us to statistically extrapolate allele frequency (Fig. 5). The colony PCR was designed to contain 3 different primers; two commercially available primers that flank that multiple cloning site of the TOPO cloning vector (M13 and T7) and one allele-specific primer that would only hybridize if the KI sequence is present. The allele specific primer for FOXA1 K295Q is *K295Q ASP RV:* 5’ GCGCCAGAGGGATCCTG 3’. The resulting PCR products would therefore produce a larger 1 kb product if the sequence is wild type/indel or a smaller 700 bp product if the KI sequence is present allowing the allele-specific primer to anneal (Fig. 5). Thus, this PCR screening method reveals the proportion of FOXA1 KI alleles within a cell line by generating unique PCR products that correlate with KI alleles, in addition to wild type/indel alleles (Fig. 5). Sanger Sequencing further validated these results and provided insight into the nature of each allele. Alternatively, a 200-300bp region surrounding the KI site may be amplified and ligated to Illumina-specific barcodes for sequenced using next-generation sequencing. The identity and ratio of different FOXA1 alleles could then be extracted using common bioinformatic platforms (34).

To further determine appropriate transcription and help estimate allele-frequency of the knockin mutants, we extracted total RNA from the knockin cell lines (Qiagen RNeasy Mini Kit), prepared Illumina-based Poly-A enriched mRNA sequencing libraries (Illumina Stranded mRNA Prep kit), and sequenced these libraries using 150 bp paired end sequencing kit to achieved at least 50 million uniquely mapped reads. The RNA-seq reads were aligned using Bowtie 1.3.0 using the default parameters, converted into Bam file using SAMTOOLS, and then loaded into Integrative Genomics Viewer (IGV) for visualization of the individual sequencing reads and the proportion of mutated nucleotides within them (Fig. 6). We were able to confirm the presence of the KI nucleotide changes as well as the silent mutations we designed to ablate the PAM recognition sites. While there were some reads that were seemingly wild-type, further investigation showed that these transcripts contained indels (insertions or deletions) immediately upstream or downstream of this region which resulted in frame shift knockouts of all wild-type proteins (Fig 6). It was also possible to estimate proportion of wild-type to KI sequences by virtue of the percentage of reads containing knockin mutations which matched the allele frequency estimations using TOPO cloning.

**Figure 6.**
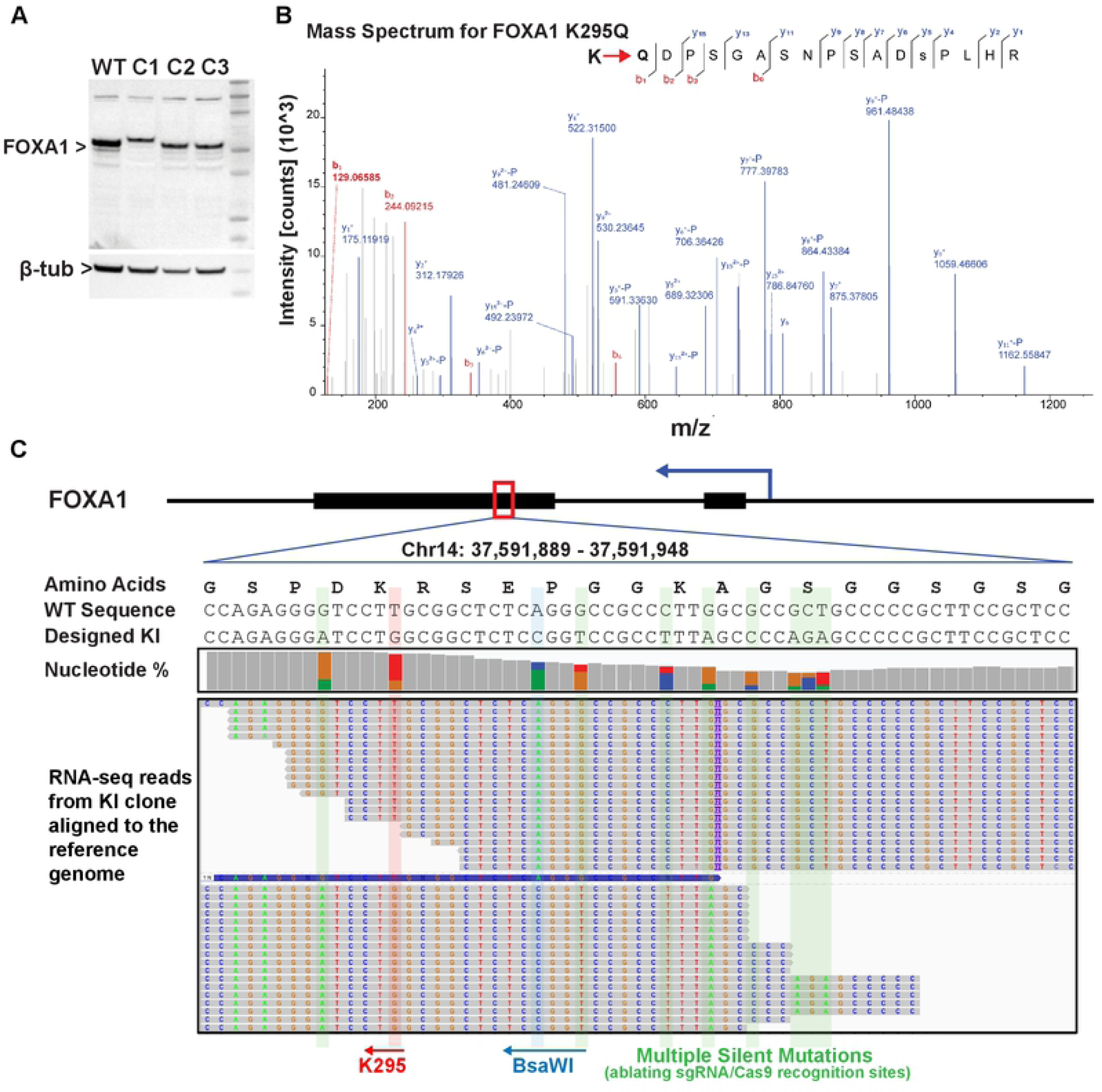
Successful replacement of wild-type FOXA1 with K295Q Knockin FOXA1 in MCF-7 cells confirmed by RNA-sequencing and mass-spectrometry. **A**. Western blot of successful FOXA1 knockin clones. Western blot showing expression of FOXA1 K295Q knockin mutants. One cell line, clone 1, shows a slightly higher molecular weight for FOXA1 potentially hinting at an insertion event. Clones 2 and 3 show appropriate molecular weight for FOXA1 with similar expression levels compared to WT FOXA1. **B**. Mass spectrum confirmation of successful K295Q mutagenesis at the endogenous locus of FOXA1. FOXA1 was immunoprecipitated from an MCF-7 knockin clone and subjected to LC-MS/MS. The fragmentation pattern confirms that lysine (K) at position 295 of FOXA1 was successfully edited to glutamine (Q). **C**. Poly-A enriched RNA-seq reads from an MCF-7 knockin clone aligned to the FOXA1 reference gene showing edited nucleotides in the mRNA transcripts of FOXA1. This gene is transcribed from the antisense strand and thus the codons are read from right to left. Taken together these data confirm successful genome editing resulting in appropriate transcription and translation of the intended knockin mutations in FOXA1.

### FOXA1 mutant protein expression analysis by western blotting and mass spectrometry

A potential consequence of editing with the CRISPR-Cas9 is repair of the resulting double strand DNA break without incorporating the repair template. Often a random number of nucleotides can be inserted or deleted at the break site and the resulting indel can lead to frameshifts in the coding sequence. In addition, it is possible that some but not all of the alleles are successfully mutated and the remaining alleles may be wild-type or knockout. These molecular events can lead to dramatic changes in protein expression or protein size, and thus, western blotting is an important quality control step in ensuring proper expression of the mutated protein (Fig. 6). Of note, it is important to ensure that the desired knockin mutations does not ablate the epitope recognized by the antibody chosen for western blot. This is especially true for monoclonal antibodies that only recognize a single epitope. Polyclonal antibodies or combinatorial uses of multiple antibodies (if available) can circumvent this issue. However, the ultimate validation of a successful amino acid substitution is via mass spectrometry. This would provide definitive evidence that the target protein is indeed expressed with the specified amino acid substitution (Fig. 6).

MCF-7 clones that contained knockin sequences based on genotyping were moved forward to western blotting. We followed standard protocols for protein extraction and western blotting to detect FOXA1 in our KI cell lines. Briefly, cells grown in a 10 cm dish were washed with ice cold PBS and collected in 1ml of fresh PBS with protease inhibitors using a cell scraper. Cells were centrifuged at 5000 rpm for 3 minutes and the remaining cell pellet was lysed with three times the pellet volume of protein lysis buffer (50 mM Tris pH8, 0.5 M NaCl, 1% NP40, 0.5% Sodium Deoxycholate, 0.1% SDS, protease inhibitors, and 50 units of Benzonase [Sigma Cat # E8263]) and incubated on ice for 30 minutes with occasional vortexing. Cell lysates were centrifuged at 12,000g for 10 minutes and the resulting supernatant was collected for western blot analysis. Protein quantification of the cell lysates was performed using Bradford protein quantification assay (Sigma Aldrich).

Protein lysates were resolved on a 4-12% gradient polyacrylamide gel, transferred to a PVDF membrane and blotted with a FOXA1 antibody (Abcam, ab23738) diluted 1:6000 in 1% BSA. A donkey anti-rabbit secondary antibody conjugated to HRP was used for visualization (GE Health, NA934-1ML) (Fig. 6).

For mass spectrometry, FOXA1 was first immunoprecipitated from MCF-7 cell lysates using a FOXA1 specific antibody (Abcam, ab23738) in order to increase the signal and remove unrelated proteins from the experiment. For immunoprecipitation (IP), protein lysates were generated from 15 cm dishes as described above and were then diluted 1:1 with dilution buffer (20 mM Tris pH8, 150mM NaCl, 2mM EDTA, 0.5% Triton X-100, and protease inhibitors) and subjected to protein quantification. For each IP assay, cell lysates containing 3000 ug total protein lysis were further diluted 1:10 using dilution buffer and incubated with 5 ug of antibody overnight at 4°C on a rocking platform. Protein A conjugated agarose beads (80 uL, 50% slurry) were washed 3 times using cold TE buffer and blocked for 2 hours using 8 uL of 10 mg/mL BSA. The blocked beads were washed 2 more times with TE buffer and transferred into the IP tube after its overnight incubation. With newly added beads, the IP tube was incubated at 4°C on a rocking platform for another 2 hours. The beads were then pelleted at 2000 rpm, 4°C for 1 minute, and washed 2 times with diluted lysis buffer (use the same dilution as for IP samples). Finally, the beads were washed another time with TE, and the protein sample was eluted in 20 uL 4X SDS loading dye and boiled at 95°C for 5 minutes. The boiled sample was centrifugated at 5000g for 3 minutes, and the supernatant was collected for gel electrophoresis and mass spectrometry analysis. The IP sample was resolved on a 4-12% pre-cast gradient gels from Invitrogen and stained using a standard Coomassie Blue staining protocol. The 50 kD band corresponding to FOXA1 was carefully excised from the gel and submitted to LC-MS/MS analysis (Fig. 6). As seen in figure 6, FOXA1 was successfully identified in the sample and the spectra for the knockin mutation (K295 to Q) is shown. Taken together, these data confirm that we have successfully edited the FOXA1 protein sequence by site-directed mutagenesis of the endogenous loci of FOXA1 in breast cancer cells.

## Discussion

### Precise editing of the endogenous loci of FOXA1 in MCF-7 cells

Completely replacing a protein coding gene sequence with one that carries targeted mutations in a polyploid cell line is inherently challenging. We have found this to be a labor-intensive process, extending over 3 months. The goal described in this protocol, to generate KI cell lines capable of producing full-length FOXA1 mutant protein, demonstrates this point well. We generated 66 clonal cell lines for our FOXA1 K295 KI experiment after antibiotic selection. Of these, 12 clones were positive for allele-specific genotyping PCR; 3 were later discarded due to apparent large indels observed via western blot validation of FOXA1 (Fig. 6). From the remaining 9 clonal cell lines, we selected 4 for TOPO cloning and sequencing. We determined small in-frame deletions within FOXA1 in 2 of these cell lines making them unusable for our further experiments. The final two cell lines contained our desired FOXA1 KI alleles along with non-functional indel alleles (knockouts). Thus, successfully replacing all wild-type versions of FOXA1 in these cells, making them ideal for downstream experiments. Based on our allele frequency estimation, we obtained 2-3 successful KI alleles of FOXA1 in our MCF-7 cell lines which contain about 3-5 alleles total of FOXA1 (Fig. 6) (33). From the RNA-seq data of these selected KI cell lines, we found that 25% to 75% mature mRNA in these cell lines carry KI mutations, and also found the presence of several KO alleles (also observed from TOPO cloning experiments). Our results suggest the KI outcome is limited by the efficiency of the HDR pathway when competing with the more facile NHEJ pathway. This scenario illustrates the success rate in generating suitable cell lines and suggests that one should aim to generate 2 or 3 times more total cell lines than actually needed. Of note, our design of the repair template used for homologous recombination allows for a second round of KI targeting and screening, which will eventually lead to complete replacement of all endogenous alleles with KI alleles.

### Determining HDR competency of targeted cell lines

Most actively proliferating cell lines are suitable for CRISPR/Cas9 directed site-specific mutagenesis. Some cell lines, however, may have inherent defects in the HDR pathway and it is important to consult existing literature for evidence of successful genome editing in the cell line of interest (27). CCLE (Broad Institute Cancer Cell Line Encyclopedia, https://portals.broadinstitute.org/ccle) is a useful resource to survey potential cell lines for defects in critical components of the HDR machinery (35). Finally, recombination activity tests may be attempted in a desired cell line for further assurance (36, 37).

### Considerations in designing targeting and repair template sequences

For reduced off-target effects, it is beneficial to choose targeting sequences that display higher specificity even if there is some compromise in efficiency. If necessary, the most likely off-target sites (as predicted by CRISPOR) can be surveyed and examined by Sanger sequencing to detect unintended alterations at these other loci.

Previous work has indicated that the DNA sequence at the Cas9 cleavage site is more likely to be repaired by an exogenous repair template than adjacent sequences (38, 39). When designing the repair template, the silent mutations (preventing repeated cleavage by sgRNA/Cas9, or to introduce restriction sites) are best introduced between the gRNA targeting sequence and the genomic editing site. These silent mutations should be clustered together, rather than dispersed, to ensure incorporation of all of the intended substitutions. Moreover, this protocol can be easily adjusted to work with Cas9 Nickase, which requires on two nearby sgRNA targeting sequences for proper recognition and cleavage which will increase on-target specificity.

## Conclusions

There are a handful of strategic paths that may be taken to further increase KI efficiency using CRISPR-Cas9 including second round of CRISPR-Cas9 KI editing using different sgRNA. Alternatively, designing a dual antibiotic selection strategy where two selectable markers are separately added to the same repair template would facilitate dual antibiotic selection for cells that integrated KI cassettes into both alleles (40). Another strategy for increasing efficiency could be chemical inhibition of the NHEJ repair pathway thus forcing the cells to engage in HDR (40-42). Finally, CRISPR-trap, an alternative genome editing strategy, could be adopted to introduce KI proteins while avoiding frameshift truncations (43, 44). The downside with this strategy, however, is the KI transcripts will lose their endogenous 3’ UTR and corresponding native gene regulation (43).

As more CRISPR editing tools emerge our ability to edit endogenous loci of polypoid cell lines will continue to increase. For example, the very recent development of CRISPR-based Prime Editing that directly writes new genetic information into a specified DNA site using a catalytically impaired Cas9 endonuclease fused to an engineered reverse transcriptase, programmed with a guide RNA that both specifies the target site and encodes the desired edit, can be imagined to readily modify the majority of endogenous proteins in polyploid cancer cell lines (45). The strategies described herein, sgRNA design, repair template design, genotype screening strategies, and allele frequency estimation, would still apply and greatly facilitate the use of these other CRISPR technologies such as prime editing.

## Declarations

### Availability of data and materials

The materials developed and the datasets analyzed during the current study are available from the corresponding author upon request.

### Author contributions

H.L.F., S.L., and J.P.G. conceived the project and developed the methodology. S.L. and C.A.T. performed the experiments with input from H.L.F and J.P.G. All authors wrote and approved the final manuscript.

## Acknowledgments

We would like to thank the members of the Franco lab and Perou lab at UNC Chapel Hill for their insightful comments and critical reading of this article.

